# Secreted Particle Information Transfer (SPIT) – A Cellular Platform for *In Vivo* Genetic Engineering

**DOI:** 10.1101/2024.01.11.575257

**Authors:** Carsten T. Charlesworth, Shota Homma, Fabian Suchy, Sicong Wang, Joydeep Bhadhury, Anais K. Amaya, Joab Camarena, Jinyu Zhang, Tze Kai Tan, Kyomi Igarishi, Hiromitsu Nakauchi

**Affiliations:** Institute for Stem Cell Biology and Regenerative Medicine, Stanford University School of Medicine, Lorry I. Lokey Stem Cell Research Building, 265 Campus Drive, Stanford, CA, USA; Department of Genetics, Stanford University School of Medicine, Stanford, CA, USA; Department of Pediatrics, Stanford University School of Medicine, Stanford, CA, USA; Department of Hematology, Stanford University School of Medicine, Stanford, CA, USA; Department of Pathology, Stanford University School of Medicine, Stanford University, Stanford, CA, USA; Division of Stem Cell Therapy, Distinguished Professor Unit, The Institute of Medical Science, The University of Tokyo, Tokyo, Japan

**Author notes:** Corresponding author: Hiromitsu Nakauchi. Co-Corresponding author: Carsten Charlesworth.

**Keywords:** *In Vivo* Gene Therapy, Human Cell Vectors, CRISPR gene editing, Regenerative Medicine, VLP, Nanomedicine Developments, Synthetic Biology, Cell Engineering, Systemic Delivery, Stem Cell Targeting

## Abstract

A multitude of tools now exist that allow us to precisely manipulate the human genome in a myriad of different ways. However, successful delivery of these tools to the cells of human patients remains a major barrier to their clinical implementation. Here we introduce a new cellular approach for *in vivo* genetic engineering, **<u>S</u>**ecreted **<u>P</u>**article **<u>I</u>**nformation **<u>T</u>**ransfer (SPIT) that utilizes human cells as delivery vectors for *in vivo* genetic engineering. We demonstrate the application of SPIT for cell-cell delivery of Cre recombinase and CRISPR-Cas9 enzymes, we show that genetic logic can be incorporated into SPIT and present the first demonstration of human cells as a delivery platform for *in vivo* genetic engineering in immunocompetent mice. We successfully applied SPIT to genetically modify multiple organs and tissue stem cells *in vivo* including the liver, spleen, intestines, peripheral blood, and bone marrow. We anticipate that by harnessing the large packaging capacity of a human cell’s nucleus, the ability of human cells to engraft into patients’ long term and the capacity of human cells for complex genetic programming, that SPIT will become a paradigm shifting approach for *in vivo* genetic engineering.

## INTRODUCTION

Two general approaches are currently being developed to apply genetic engineering tools to patients. The first are *ex vivo* genetic engineering approaches, where patient cells are isolated from the body, genetically engineered *ex vivo* and then transplanted back into the body^4, 5, 6^. Although this approach is clinically efficacious, its application is restricted to cell types that are amenable to this *ex vivo* process, primarily hematopoietic cell types. The second approach is to deliver genetic engineering technologies to cells directly *in vivo* through the use of recombinant viral vectors such as adeno associated virus (AAV, carrying up to ∼4.5kb of DNA) or chemically defined platforms such as lipid nanoparticles (LNPs, have delivered up to ∼10-20kb of RNA *in vivo*)^7, 8^. However, the successful clinical application of these *in vivo* technologies has primarily been restricted to the liver and current platforms are limited in the amount of genetic information they can deliver to cells, typically limited to a single gene due to packaging limitations^9, 10^.

Compared to the limited packaging capacities of contemporary *in vivo* gene therapy delivery platforms, a human cell’s nucleus contains approximately 6 billion base pairs of information^11^. We thus postulated that if we could use human cells as vectors for *in vivo* gene therapy that we could vastly increase the amount of genetic information we could deliver and thus manipulate *in vivo*. We hypothesized that human cells could be applied as vectors for *in vivo* gene therapy by modifying them to **<u>S</u>**ecrete a genetic engineering enzyme within a **<u>P</u>**article, that **<u>T</u>**ransfers this enzyme into a recipient cell, where it manipulates genetic **<u>I</u>**nformation. We term this cellular *in vivo* gene therapy approach **<u>S</u>**ecreted **<u>P</u>**article **I**nformation **<u>T</u>**ransfer (SPIT). Here we successfully demonstrate the application of SPIT for cell-cell delivery of Cre and the CRISPR-Cas9 system for genetic engineering, show that genetic logic can be incorporated to regulate SPIT and successfully present the first application of human cells as a delivery platform for *in vivo* genetic engineering, in immunocompetent mice.

## RESULTS

### Identifying a delivery modality to achieve SPIT

To successfully demonstrate proof-of-concept for SPIT, we first needed to identify a delivery modality for genetic engineering that could both be secreted by a cell and deliver genetic engineering enzymes to a cell. We identified Viral and Vesicle (VLPs) like particles as modalities that met these requirements^12^. Evaluating a commercially available VLP platform (gesicles) for its ability to deliver Cre to Ai14 reporter cells, via tdTomato expression^13^. We found that VLPs could effectively deliver Cre to multiple primary cell types *in vitro* including mouse fibroblasts and mouse hematopoietic stem cells (HSCs) (**Figure 1A-C, Supplemental Figure 1 A-C**).

**Figure 1:**
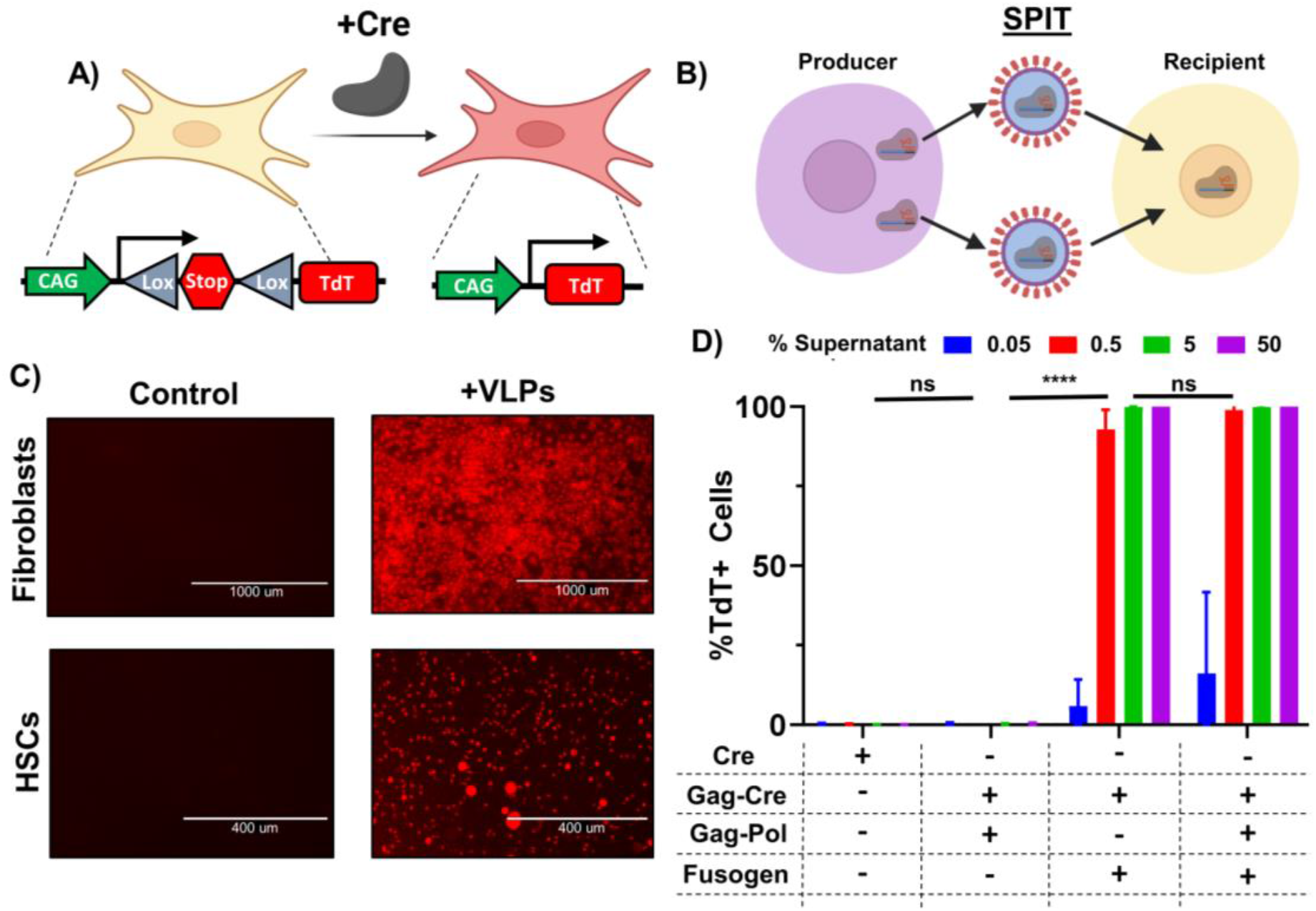
Retroviral VLPs Are an Efficacious Modality to Accomplish SPIT. **A)** Schematic of the Ai9/Ai14 reporter system used to detect delivery of Cre to cells via expression of tdTomato (TdT). **B)** Schematic showing the general concept of SPIT for facilitating cell-cell delivery of genetic engineering technologies. **C)** Representative images of tdTomato expression in fibroblasts and HSCs 3 days post application of VLPs (gesicles) compared to untreated (control) cells. **D)** Bar graph showing the frequency of TdT+ Ai14 fibroblasts following the application of different VLP formulations to cells, analyzed for TdT expression 3 days post application of VLPs by FACs (ANOVA, n=3, mean± s.d). P<0.0001 = ****.

Having identified VLPs as a potentially efficacious technology to accomplish SPIT with, we next sought to identify a VLP system that had the following features: (1) was able to deliver CRISPR-Cas ribonucleoproteins (RNP) for genome editing, (2) could package this RNP without the use of any kind of synthetic chemical dimerization, and (3) required the use of less than 2/3rds of a viral genome for its function, in order to follow NIH guidelines regarding the introduction of viral genes into eukaryotic organisms *in vivo*^*14*^. We identified the murine leukemia virus (MLV) retroviral VLP platform, Nanoblades, as one that closely met most of these criteria^15^. This system facilitates the packaging and delivery of a protein of interest (POI) (including a CRISPR-Cas RNP) to cells, through its direct fusion to the C-terminus of retroviral Gag. However, VLPs from this system are produced using more than 2/3rds of a viral genome (**Supplemental Figure 1E**).

To adapt this VLP platform for a proof-of-concept demonstration of SPIT *in vivo*, we screened to determine if any of the viral genes used to produce these VLPs could be eliminated, without impairing their ability to deliver Cre to Ai14 reporter fibroblasts *in vitro* (**Figure 1A**). We found that while elimination of the fusogen (VSV-g) from VLPs completely abolished their efficacy, GAG-pro-pol could be eliminated from VLPs without any statistically significant effect on the delivery of Cre to reporter cells (**Figure 1D**) (P<0.0001, ANOVA, n=3). These results demonstrate that despite the role of Gag-pro-pol in proteolytically releasing a POI from Gag during VLP formation, it was not essential for the packaging, delivery, or activity of a POI by VLPs (**Supplemental Figure 1D-E**)^15^. In addition, by eliminating Gag-Pro-Pol from VLPs and only producing VLPs through the use of Gag-Cre and VSV-g, we could produce functional VLPs using less than 2/3rds of a viral genome making this VLP formulation suitable for application as part of a SPIT platform *in vivo*.

### Development of a VLP-SPIT platform

Having identified a VLP conformation that met all our requirements, we next determined its efficacy for achieving SPIT *in vitro*. Gag-Cre and VSV-g were transfected into wild type 293T cells (wt293T) and these cells were then mixed 1-day post-transfection with reporter Ai9 293T cells at a ratio of 1:1. Tracking Cre recombination in reporter cells over time via tdTomato expression, we found that this VLP-SPIT approach could successfully be applied *in vitro* to facilitate cell-cell genetic engineering. An average of 14% of cells became tdTomato positive after 6 days of co-culture with SPIT cells, compared to 0.1% when reporter cells were cultured alone (P<0.0001 at day 6, ANOVA, n=3) (**Figure 2A-C**). Performing the same experiment using primary mouse fibroblasts, we found that SPIT could also be achieved in primary cells. With a mean of 0.79% of cells becoming positive for tdTomato expression after 4 days of co-culture, compared to 0.02% when reporter cells were cultured alone (ANOVA, P=0.0013, n=3) (**Supplemental Figure 3A**).

**Figure 2:**
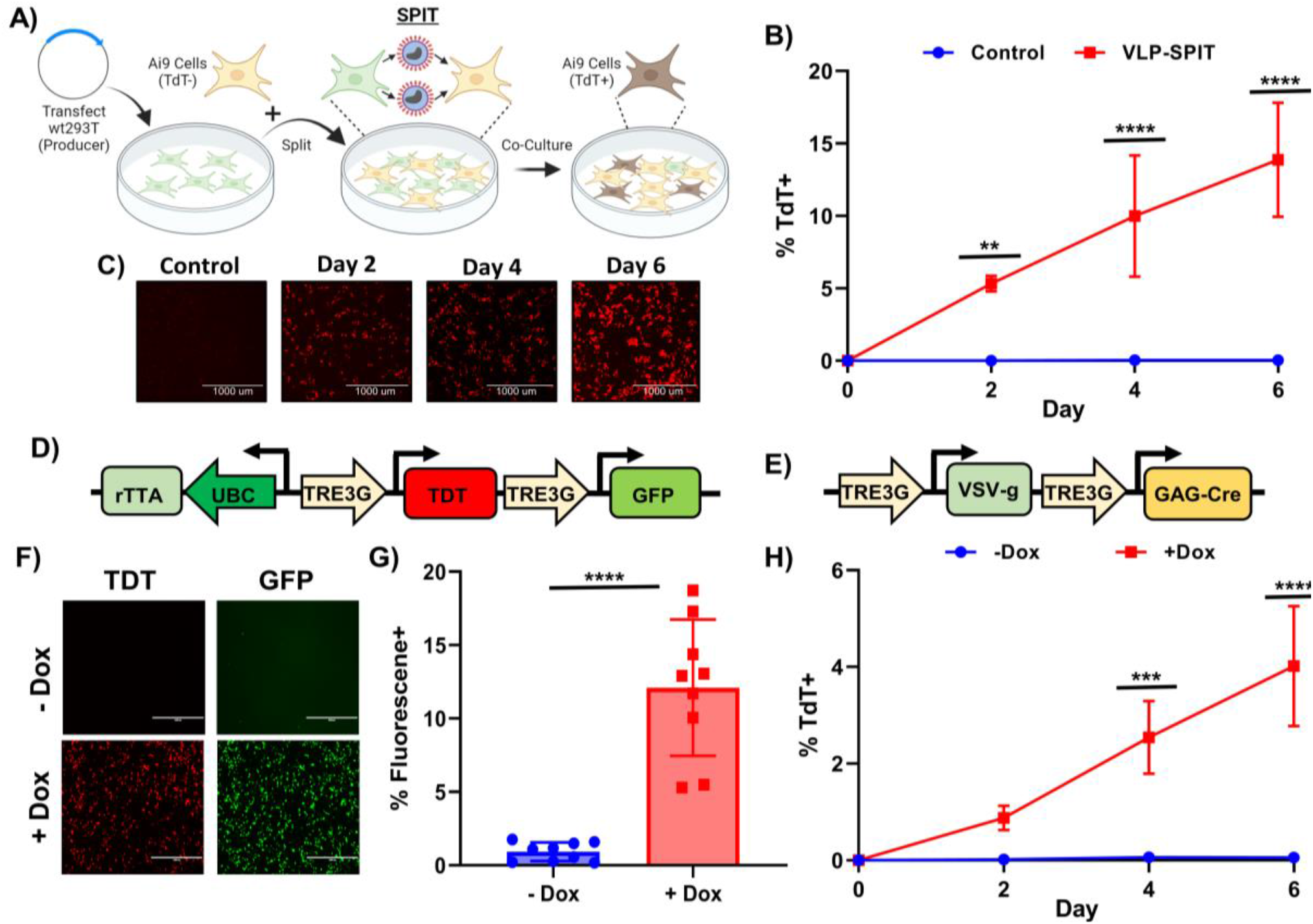
VLPs Facilitate SPIT *In Vitro* and Regulation of SPIT Using Genetic Logic. **A)** Schematic outlining the experimental procedure for testing VLP-SPIT *in vitro*. **B)** Line graph showing the frequency of tdTomato+ cells over time when attempting VLP-SPIT *in vitro* using 293T cells compared to the culture of reporter cells alone (ANOVA, n=3, mean± s.d). **C)** Representative images from fluorescence microscopy of tdTomato expression in cells from experiments performed in figure 2B. **D)** Design of an all-in-one doxycycline inducible vector. **E)** Representative images from fluorescence microscopy of GFP and tdTomato fluorescence in cells transfected with the vector shown in figure 2D when doxycycline was present or absent in media. **F)** Bar graph showing the total percentage of cells that were positive for GFP or tdTomato expression as determined by flow cytometry when doxycycline was present or absent in the media after transfection of all-in-one doxycycline inducible vectors, cells were analyzed one day after transfection (n = 9, t-test, mean± s.d) **G)** Design of an all-in-one doxycycline inducible VLP-SPIT construct. **H)** Line graph showing the frequency of tdTomato+ cells in culture over time when Ai9 reporter 293T cells are co-cultured with 293T cells transfected with the vector shown in figure 2G, when doxycycline is present or absent in the media. (ANOVA, n=3, mean± s.d). *** = P < 0.005, **** = P < 0.0005. Dox = Doxycycline, TdT = tdTomato, GFP = Green Fluorescent Protein.

We next sought to demonstrate the potential for regulating the activity of SPIT using genetic logic, by developing a SPIT vector where cell-cell delivery of a genetic engineering enzyme only occurred when a small molecule was applied to cells. We first constructed an all-in-one inducible vector, where the expression of desired genes was regulated by the addition or withdrawal of doxycycline^16^. Several different vectors were designed with different promoter/gene orientations, and the efficacy of these vectors for doxycycline regulatable gene expression in cells determined using tdTomato and GFP (**Figure 2D & 2F-G, Supplemental Figure 2**). Gag-Cre and VSV-g were then placed under the control of doxycycline inducible promoters within the most efficacious all-in-one construct (Vector A) (**Figure 2E**, Supplemental **Figure 2C**). This inducible SPIT construct was then transfected into wt293T cells and these cells co-cultured with Ai9 293T cells in the presence or absence of doxycycline. Comparing the two conditions after 6 days of co-culture we found an average of 4% of cells were positive for tdTomato expression when cells were co-cultured in the presence of doxycycline, compared to only 0.1% when cells were co-cultured in its absence (ANOVA, P<0.0001, n=3) (**Figure 2H**). These results demonstrate proof-of-principal for the incorporation of genetic logic into SPIT.

**Figure 3:**
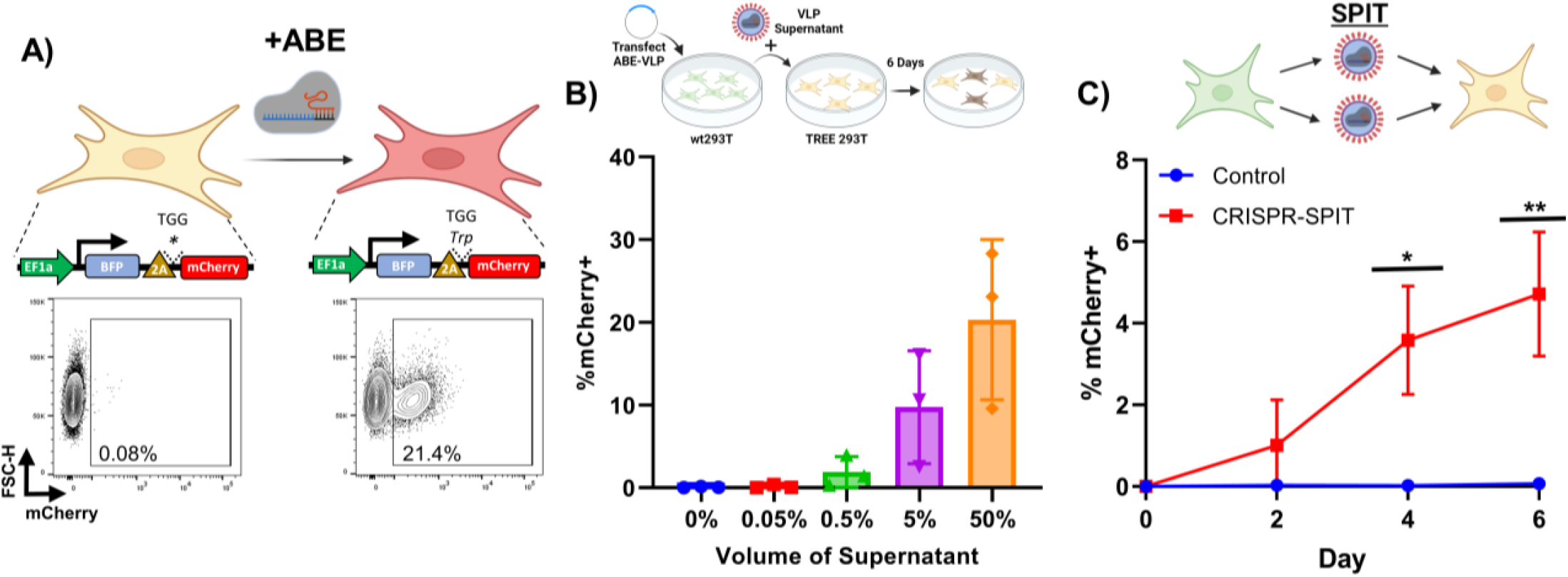
SPIT Can Deliver CRISPR-Cas RNPs for Genome Editing. **A)** (Top) Schematic of the TREE reporter system used to assess if adenine base editors were successfully delivered to receiver cells, (Bottom) representative FACs plot of mCherry expression in unedited TREE 293T cells (left) and edited TREE 293T cells (right). **B)** (Top) Schematic of the experiment performed to test CRISPR VLPs. (Bottom) Bar graph showing the frequency of mCherry+ TREE reporter 293T cells, when ABE VLPs were produced in 293T cells and the supernatant from these producer cells was applied to reporter cells across a dose titration (n=3, mean± s.d). **C)** (Top) Schematic of the experiment performed to test CRISPR-SPIT. (Bottom) Line graph showing the results from CRISPR-SPIT experiments. CRISPR-SPIT constructs were transfected into 293T cells, which were then collected and co-cultured at a 1:1 ratio with TREE reporter 293T cells (ANOVA, n=3, mean± s.d). * = P < 0.05, ** = P < 0.005. ABE = CRISPR-Cas9 adenine base editor, VLP = Viral like particles, wt293T = wild type 293T cells.

### CRISPR-SPIT

To determine if SPIT could also be applied to deliver CRISPR-Cas RNPs for gene editing, we set up a reporter cell line by introducing a previously described transient reporter for editing enrichment (TREE) into the genome of 293T cells. Where successful gene editing of cells by a CRISPR-Cas9 adenine base editor (ABE) could be detected via the expression of mCherry (**Figure 3A, Supplemental Figure 3B**)^17^. Using this reporter system, we found that we could successfully produce VLPs that could deliver an ABE RNP for gene editing (**Figure 3B**). Notably while Gag-pro-pol was not necessary for achieving gene editing in reporter cells with ABE VLPs, its inclusion in the VLP production led to a statistically significant improvement in the rate of gene editing in reporter cells. Due to the presence of a 3x nuclear export signal that is removed from ABE during VLP formation and regulates the cellular localization of ABE and Gag in producer versus recipient cells (ANOVA, P = 0.0011, n=2-3) (**Supplemental Figure 3C-D**)^18^. Transfecting ABE VLP plasmids into wt293T cells and then co-culturing these cells with TREE 293T reporter cells at a ratio of 1:1, we found that SPIT could also be applied to deliver ABE for cell-cell genetic engineering. An average of 4.7% of mCherry positive cells were observed 6-days post co-culture, compared to 0.1% when reporter cells were cultured alone (ANOVA, P<0.0001, n=3) (**Figure 3C**).

### SPIT enables in vivo genetic engineering

Having established that we could apply SPIT for cell-cell genetic engineering *in vitro*, we next sought to demonstrate proof-of-concept for the application of SPIT *in vivo*. Two cell types that were easily amenable to genetic modification by plasmid transfection: C57BL6/j mouse embryonic fibroblasts (MEFs) and human 293T cells, were selected as potential vectors for SPIT. The efficacy of each cell type as a SPIT vector was then compared through transfection of a luciferase expression plasmid into cells, followed by intra-peritoneal (IP) injection of 2e7 transfected cells into mice. Tracking luciferase expression over time, we found that both cell types could transiently engraft and express a transfected gene for at least 2-days *in vivo (***Supplemental Figure 4**). No statistically significant differences in the persistence of luciferase expression were found between the cell types transplanted or the immunological setting of the host they were transplanted into (xenogeneic, allogeneic, or syngeneic hosts) (ANOVA, n=2-3).

Finding that human 293T cells could transiently persist and express transfected genes when transplanted into immunocompetent mice, we proceeded to utilize these cells as vectors for SPIT *in vivo*. SPIT 293T cells were generated by transfection of Gag-Cre and VSV-g into cells and 2e8 of these SPIT 293T cells were then IP injected into Ai14 reporter mice (**Figure 4A**). After 1.5 weeks, mice were euthanized and Cre recombination determined by tdTomato expression. Dissection of mice revealed clear tdTomato expression in multiple organs and tissues of SPIT-treated mice including: the diaphragm, liver, spleen, perigonadal fat, and in some cases the intestines (**Supplemental Fig 5A**). For a quantitative analysis of Cre-mediated recombination, solid organs were extracted from mice and the intensity of tdTomato fluorescence in each organ measured with an IVIS imager. Statistical analysis of the fluorescence intensity of each organ indicated a significantly higher amount of tdTomato fluorescence in the livers (mean ROI/Background control = 1.3, SPIT = 4, p = 0.025, n = 3-5) and spleens (mean ROI/Background control = 0.88, SPIT = 2, p = 0.003) of SPIT-treated mice compared to untreated controls (**Figure 5B-C**). Notably, the intestines of some SPIT-treated mice also showed significantly increased tdTomato expression compared to controls by IVIS imaging, albeit with considerable sample to sample variability.

**Figure 4:**
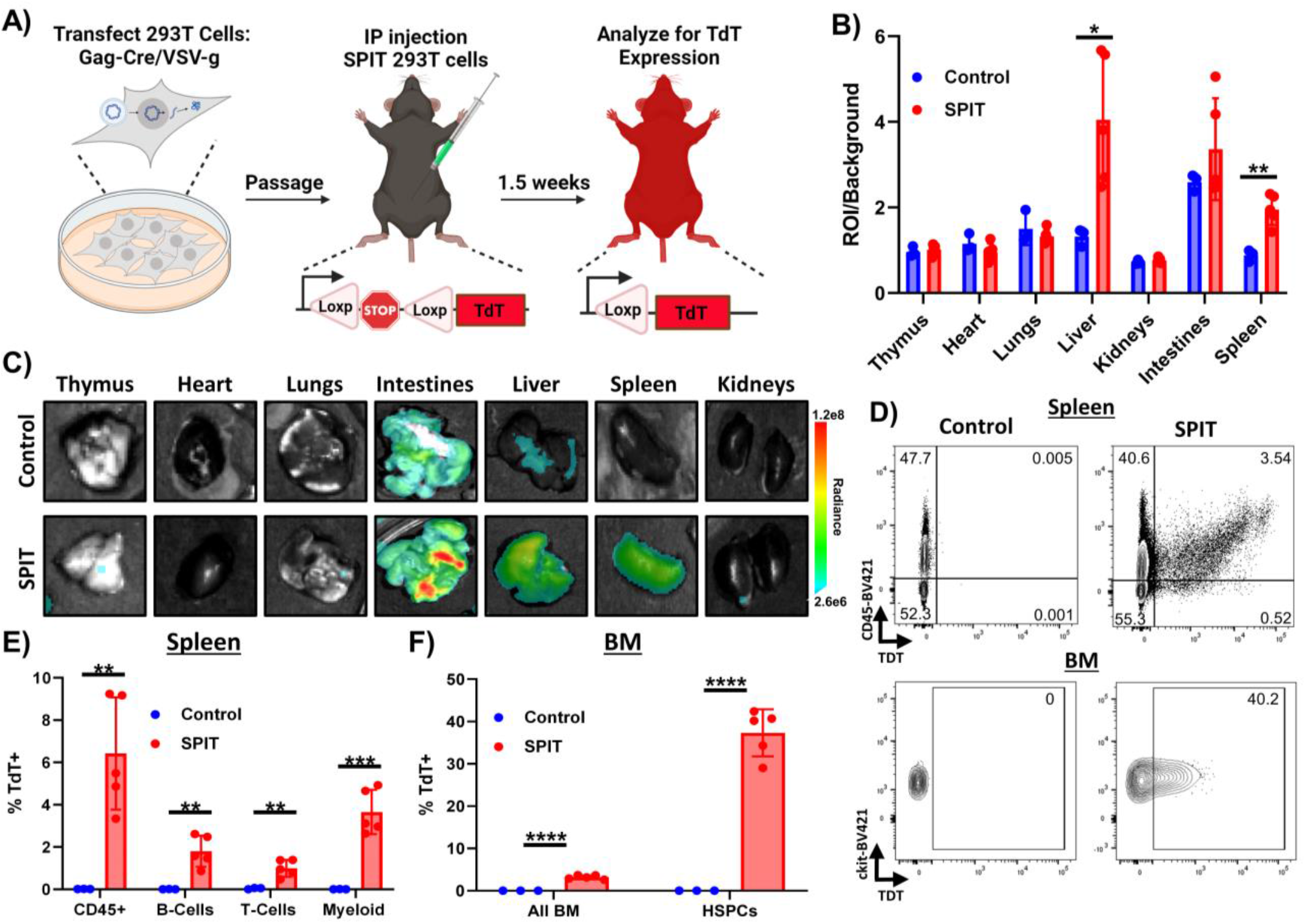
*In Vivo* Genetic Modification of Immunocompetent Mice Via SPIT. **A)** Schematic outlining the *in vivo* experiment performed and analysis. **B)** Bar graph showing the total photons/second of different organs (ROI) from experimental and control mice imaged for tdTomato fluorescence using IVIS, divided by background (t-test, n = 3-5, mean± s.d). **C)** Representative images of organs analyzed for tdTomato expression by IVIS. **D)** Representative FACs plots from flow cytometry of untreated and SPIT treated Ai14 mice analyzing for tdTomato expression. Top panels are representative FACs plots analyzing cells from the spleen while bottom panels are representative FACs plots from analyzing HSPCs (LSK) in the bone marrow. **E)** Bar graph showing the frequency of tdTomato positive hematopoietic cells in the spleen of mice treated with SPIT versus untreated control Ai14 mice, as determined by flow cytometry (t-test, n=3-5, mean± s.d). **F)** Bar graph showing the frequency of tdTomato positive cells in the BM as determined by flow cytometry (t-test, n=3-5, mean± s.d). BM = Bone Marrow, HSPCs = Hematopoietic Stem and Progenitor Cells, ROI = Region of interest, Radiance = photons/s. * = P < 0.05, ** = P < 0.005, *** = P < 0.0005, **** = P < 0.00005.

To ascertain whether SPIT delivered Cre recombinase systemically and to validate Cre recombination at a single cell level, cells from the spleen, peripheral blood (PB), and bone marrow (BM) of mice were collected for flow cytometric analysis. In the spleen, the 1.2-fold increase in fluorescence intensity measured by IVIS in SPIT-treated mice corresponded to 6.4% of CD45^+^ splenocytes expressing tdTomato (Cre-transduced cells), compared to 0.02% of splenocytes from the control (**Figure 4D-E, Supplemental Figure 6**). Further examination of specific hematopoietic lineages within the spleen found that 1.8% of B-cells (CD45^+^CD45R^+^), 1% of T-cells (CD45^+^CD4/8^+^), and 3.7% of Myeloid Cells (CD45^+^CD11b/Ly6G^+^) in the spleen were positive for tdTomato expression in SPIT-treated mice, compared to less than 0.1% of cells in controls.

PB analyses of SPIT-treated mice revealed significantly lower levels of Cre recombination in circulating mononuclear cells than in the spleen. An average of 0.04% of CD45^+^ PB cells were positive for tdTomato expression in SPIT-treated mice; nonetheless tdTomato expression could clearly be detected (**Supplemental Figure 5B & 6**). In contrast, significantly higher levels of recombination could be detected in cells within the BM, with an average of 2.9% of BM cells positive for tdTomato expression in SPIT-treated mice, compared with 0.03% of cells in the control group. Interestingly, a significantly higher proportion of hematopoietic stem and progenitor cells (HSPCs; Lineage^-^cKit^+^Sca1^+^; LSK) were positive for tdTomato expression compared to the general population of cells in the BM, with an average of 37% of HSPCs in SPIT-treated mice positive for tdTomato expression, compared to 0% of cells in control mice (**Figure 4D & F, Supplemental Figure 7**). No differences in the frequency of different cell types could be detected between SPIT-treated and control mice in any of the organs analyzed by flow cytometry (**Supplemental Figure 5C-E**). Collectively, these results convincingly demonstrate that human cells are capable of serving as vectors *in vivo* delivery of genetic engineering enzymes via SPIT. Achieving local and systemic delivery within an immunocompetent setting, including to adult stem cells.

## DISCUSSION

Three facets of human cells make them ideal delivery vectors for *in vivo* genetic engineering: 1) the amount of genetic information they can incorporate, 2) their capacity to perform complex genetic/cellular logic, and 3) their ability to engraft into patients’ long term. Here through the use of VLP technology we successfully developed an approach to apply human cells as vectors for *in vivo* genetic engineering by Secreted Particle Information Transfer or SPIT. We showcase the versatility of SPIT by employing it to deliver both Cre recombinase and a CRISPR-Cas9 adenine base editor for precision genome engineering. Additionally, we illustrate the integration of genetic logic within the SPIT platform and present the first demonstration of human cells as vectors for *in vivo* genetic engineering. Our findings underscore the immense promise that human cells have as vectors for the delivery of a diverse array of genetic engineering tools *in vivo*, including transcription factors (such as OKSM), telomerases, zinc fingers, TALENs and CRISPR-Cas systems, among others.

The use of human cells for *in vivo* genetic engineering opens up new avenues and potential approaches for *in vivo* gene therapy. The vast and virtually limitless payload capacity of a human cell’s nucleus has the potential to enable multiplexed *in vivo* genetic engineering on a scale that is currently unattainable using conventional delivery platforms. While the ability of human cells to engraft into patients’ long-term means that SPIT can be applied to deliver genetic engineering enzymes to a patient continuously, for an indefinite amount of time. We also highlight how the innate ability of cells for intricate genetic programming can be harnessed to create a chemically regulated SPIT platform, offering a glimpse into future applications that could marry SPIT with genetic/cellular logic for gene therapy. For instance, SPIT could be incorporated into a CAR-T cell therapy for the genetic treatment of cancer.

Central to our focus was the development of a SPIT platform that could deliver a CRISPR-Cas RNP. This is due to the transformative nature of these systems for genetic engineering and the challenge that cellular RNAses pose to the stability of an sgRNA. Unlike mRNA delivery systems that require chemical modification of an sgRNA to achieve gene editing, a Cas protein can protect an sgRNA from cellular RNAses when bound to it and delivered to cells as an RNP^19^. While a number of different VLP systems have been developed that can deliver mRNA to cells, these VLPs have struggled to achieve gene editing with CRISPR-Cas systems unless an sgRNA is supplied to cells exogenously^12, 20^. Here we circumvented this challenge to achieve cell-cell genetic engineering with a CRISPR-Cas9 adenine base editor, by SPIT, via its packaging and delivery to recipient cells as an RNP.

Further work will be needed to transition SPIT from proof-of-concept to a clinically translatable therapy. Although we have successfully adapted retroviral VLPs for SPIT, the clinical potential of this strategy is constrained by factors such as the cytotoxicity of the fusogen used (VSV-g), the indiscriminate targeting of cell types by SPIT, and the immunogenicity of viral components as well as gene editing enzymes themselves^21, 22^. Future Optimizations of SPIT will include the exploration of non-viral, synthetic, and humanized VLP alternatives, enhancing target specificity to reduce off-target effects, and using non-transformed donor cells as vectors for delivery.

In summary, our work not only establishes human cells as a novel delivery platform for *in vivo* genetic engineering but also showcases the inherent advantages of such a system— namely the substantial amount of genetic information human cells can package, their capacity for extended delivery of genetic engineering enzymes to cells *in vivo* and their compatibility with complex genetic programming. While significant further work and careful consideration is required to clinically translate SPIT, the versatility and adaptability of this approach suggests that it could become a transformative tool in modern medicine.

## MATERIALS AND METHODS

### Data reporting

No statistical methods were used to predetermine sample size. The experiments were not randomized, and the investigators were not blinded to outcome assessment.

### Plasmids

Plasmids GAG-Cre (119971), GAG-Pro-Pol (35614), VSV-g (12259), GAG-ABE (181751), ABE (164415), and the sgRNA for the TREE reporter system (164413) were all obtained from Addgene. To generate Ai9 reporter 293T cells, the reporter sequence from the Ai9 plasmid (22799) was cloned in between the homology arms of an AAVS1 HDR construct from addgene (64215). To generate TREE 293T reporter cells, the ABE TREE reporter construct (164411) was cloned from its original vector into a piggybac plasmid with a puromycin selection cassette (pb-TREE). The overall function of the reporter was not altered, however the fluorescent reporter activated by ABE gene editing was swapped from GFP to mCherry during cloning. Doxycycline inducible elements were synthesized by gene universal from previously published sequences and different doxycycline inducible plasmids were then constructed from this synthesized vector using Gibson assembly. The luciferase expression vector was cloned in-house and the luciferase gene a gift from Dr. Atsushi Miyawaki (PMID: 29472486), while the hyperbase plasmid used to generate reporter cell lines by Piggybac insertion was a gift to the Nakauchi lab from Dr. Yasuhide Ohinata (PMID: 24667806).

### VLP production and Screening

For experiments where VLPs were isolated from the supernatant of producer cells and applied to recipient cells the following procedures were followed. 7e6 293T cells were plated onto a 10cm dish, one day after plating 2ug of each plasmid (maximum 8 ug of plasmid) was transfected into cells using 60 ug of PEI max (Polysciences). Producer cells were then maintained for 3 days after transfection. After which, the supernatant was taken from cells and spun down once at 2000g for 10 minutes. It was then run through a 0.45 um filter and concentrated using an Amicon 100kDa filter (Millipore Sigma). Reporter cells were prepared on the same day that supernatant was collected from producer cells and cultured on a 48 well plate at a concentration of 10,000 cells/cm^2. After plating the reporter cells, concentrated VLPs were applied across a dose titration. For experiments where VLPs delivered Cre to Ai14 fibroblasts, reporter cells were analyzed for tdTomato expression 3 days after the application of VLPs by flow cytometry. In cases where VLPs delivered ABE8, reporter cells were split 3 days after the application of VLPs and cultured for an additional 3 days, after which cells were analyzed for mCherry expression by flow cytometry.

### SPIT Experiments and Transfections

Both 293T cells and mouse fibroblasts were cultured in DMEM (Gibco) with 1x sodium pyruvate (Gibco), 1x glutamax (Gibco), 1x penicillin/streptomycin (Gibco), 1x non-essential amino acids (Gibco), and 10% FBS (Gibco). For experiments where 293T cells were co-cultured to demonstrate SPIT, 293T cells were plated at a concentration of 9e4 cells/cm^2 one day before transfection on a 12 well plate. Cells were then transfected with 500ng of plasmid or 100ng of plasmid in the case of VSV-g. Plasmids were prepared in optimem (Gibco) and transfected using 6 ug of PEI max. Cells were collected 1 day after transfection and then co-cultured with reporter 293T cells at a ratio of 1:1 in 12 well plates at a concentration of 90,000 cells/cm^2. Cells were split and analyzed by FACs once every two days post co-culture and split 1:1. For any other experiment where plasmids were transfected into 293T cells, unless otherwise specified, the same chemical transfection procedure was used. In some cases where doxycycline inducible constructs were tested for chemical regulation of fluorescent proteins charcoal stripped FBS was used in cell media. For SPIT experiments with primary fibroblasts, C57BL6 MEFs (ATCC) were plated at 20,000 cells/cm^2 on a 6 well dish. 24 hours after plating cells were transfected with GAG-Cre (800ng) and VSV-g (100ng) using lipofectamine 3000 (Invitrogen) following the manufacturers’ instructions. One day later cells were collected and then co-cultured with Ai14 fibroblasts at a ratio of 1:1 on six well plates at a concentration of 20,000 cells/cm^2 and passaged/analyzed by flow cytometry once every two days thereafter. For experiments where a doxycycline inducible plasmid was used, doxycycline was added to cells at a concentration of 2ug/ml (Sigma-Aldrich).

### Reporter Cell Lines

Ai14 fibroblasts and mouse HSCs were generated from Ai14 mice as previously described^23, 24, 25^. Ai9 293T cells were generated through the knock in of the Ai9 reporter construct into the AAVS1 locus by homology dependent repair and puromycin selection (2 ug/ml). TREE 293T reporter cells were generated by piggyBAC insertion of the pb-TREE reporter construct into cells by chemical transfection of the pb-TREE plasmid together with the hyperbase plasmid. Reporter cells were then selected with puromycin (2 ug/ml) (Sigma-Aldrich).

### Transplantation of Cells into Mice

Ai14 (007914) mice were either purchased from Jackson laboratories or bred in-house, while CD1 (022) mice were purchased from Charles River Laboratories. For experiments tracking luciferase expression *in vivo* 1.5e7 293T cells or 2.9e6 C57BL6/j MEFs (ATCC) were plated onto 15cm plates one day before transfection. Cells were then transfected one day after plating with 10ug of our luciferase expression vector using 60ug of PEI max per plate. Twelve hours post transfection cells were collected from plates using Tryple (Gibco), 2e7 cells were re-suspended in 200 ul of PBS and injected intra-peritoneally into mice using a 22-gauge needle. For *in vivo* SPIT experiments 6.5e6 293T cells were plated onto a 15cm plate two days before transfection. Two days post plating each plate was transfected with 9 ug of GAG-Cre and 1 ug of VSV-g, using 60 ug of PEI max per transfection. Twelve hours post transfection cells were collected and 2e8 cells were re-suspended in 400ul of PBS and injected into Ai14 reporter mice intra-peritoneally using a 22-gauge needle. All animal protocols were approved by the Administrative Panel on Laboratory Animal Care at Stanford University.

### Imaging

Pictures of cells in culture were taken using an EVOS FL imager. Following dissection of SPIT-treated mice images of the peritoneum were taken using Xcite-GR fluorescence flashlight (510-540nm) and filter (600nm longpass) (Nightsea). For quantitative imaging of organs, a Lago IVIS imager (SI imaging) was used with an excitation of 535nm and emission of 590nm. For luciferase imaging experiments mice were injected with 0.15 mmol of Akaluciferin (gift from Kuragani Kasei), after ten minutes spectral luminescence from mice was measured using an IVIS imager. The amount of radiance from each organ or mouse was determined using Aura imaging software (SI imaging).

### Flow cytometry

For all experiments that were performed *in vitro* cells were analyzed using a LSR fortessa flow cytometer (BD). Cells from the spleen, peripheral blood and bone marrow were isolated and prepared for flow cytometry as previously described, bone marrow was collected from the femurs of mice. Splenocytes and peripheral blood cells were stained with the following antibodies for 30 minutes at 4 C: FITC CD11b (M1/70; eBioscience), FITC-GR1/Ly-6G (1A8; BioLegend), APC-CD4 (RM4-5; Invitrogen), APC-CD8 (53-6.7; Invitrogen), APC Efluor 780-B220 (RA3-6B2; Invitrogen), BV421-CD45.2 (104; Invitrogen). Cells from the bone marrow were first stained with the following combination of biotinylated antibodies (lineage cocktail) for 30 minutes at 4 C: Gr-1 (RB6-8C5; Biolegend), Ter-119 (TER-119; Invitrogen), CD4 (RM4-5; Biolegend), CD8 (54-6.7; BioLegend), B220 (RA3-6B2; Biolegend), IL-7R (A7R34; Biolegend). After which cells were washed with PBS and then stained in the following cocktail for 30 minutes at 4 C: BV421-ckit (2B8; Biolegend), FITC-Sca1 (D7; Biolegend), APC/Efluor780-streptavidin (Biolegend). After staining with antibodies cells were re-suspended in PBS with propidium iodide at a concentration of 1 ug/ml, after which cells were analyzed using a FACs Aria flow cytometer (BD).

### Statistical Analysis

two-way ANOVA tests and unpaired two-tailed t-tests were performed as indicated in the figures, using Prism 9 software.

## ACKNOWLEDGEMENTS

We thank A. Wilkinson and S. Vaidyanathan for their advice, and assistance in editing this manuscript. This research was funded by Stinehart-Reed, 2020-ISCBRM internal grants. C.T.C. is supported by the National Science Foundation Graduate Research Fellowship under Grant No. (DGE-1656518). T.K.T acknowledgements funding from A*STAR National Science Scholarship. A.K.A was funded by Elizabeth Nash Memorial Fellowship from CFRI and the Stanford Maternal and Child Health Research Institute Postdoctoral Support. J.B. was supported by the International Postdoc grant from the Swedish Research Council (2017-00344) and the Assar Gabrielsson Foundation, Sweden. H.N was supported by the NIH (R01DK116944; R01HL147124), the Ludwig Foundation, the Stinehart-Reed Foundation, and the Japan Society of the Promotion of Science. ChatGPT was responsibly used for editing this text and for generating cover art.

## AUTHOR CONTRIBUTIONS

C.T.C conceptualized the research, performed experiments, analyzed data and prepared the manuscript. S.H, conceptualized elements of experiments and performed experiments. S.W, F.S, J.B, A.K.A, J.Z, KI and A.T performed experiments and edited the manuscript. J.C helped conceptualize the research and edit the manuscript. H.N supervised the research, helped conceptualize experiments and edited the manuscript.

## STATEMENT ON USE OF GENERATIVE AI

During the preparation of this work the authors used ChatGPT in order to for editing of the manuscript. After using this tool/service, the authors reviewed and edited the content as needed and take full responsibility for the content of the publication.

