## Supplemental Figures for "Secreted Particle Information Transfer (SPIT) – A Cellular Platform for *In Vivo* Genetic Engineering"

### Supplementary Figures

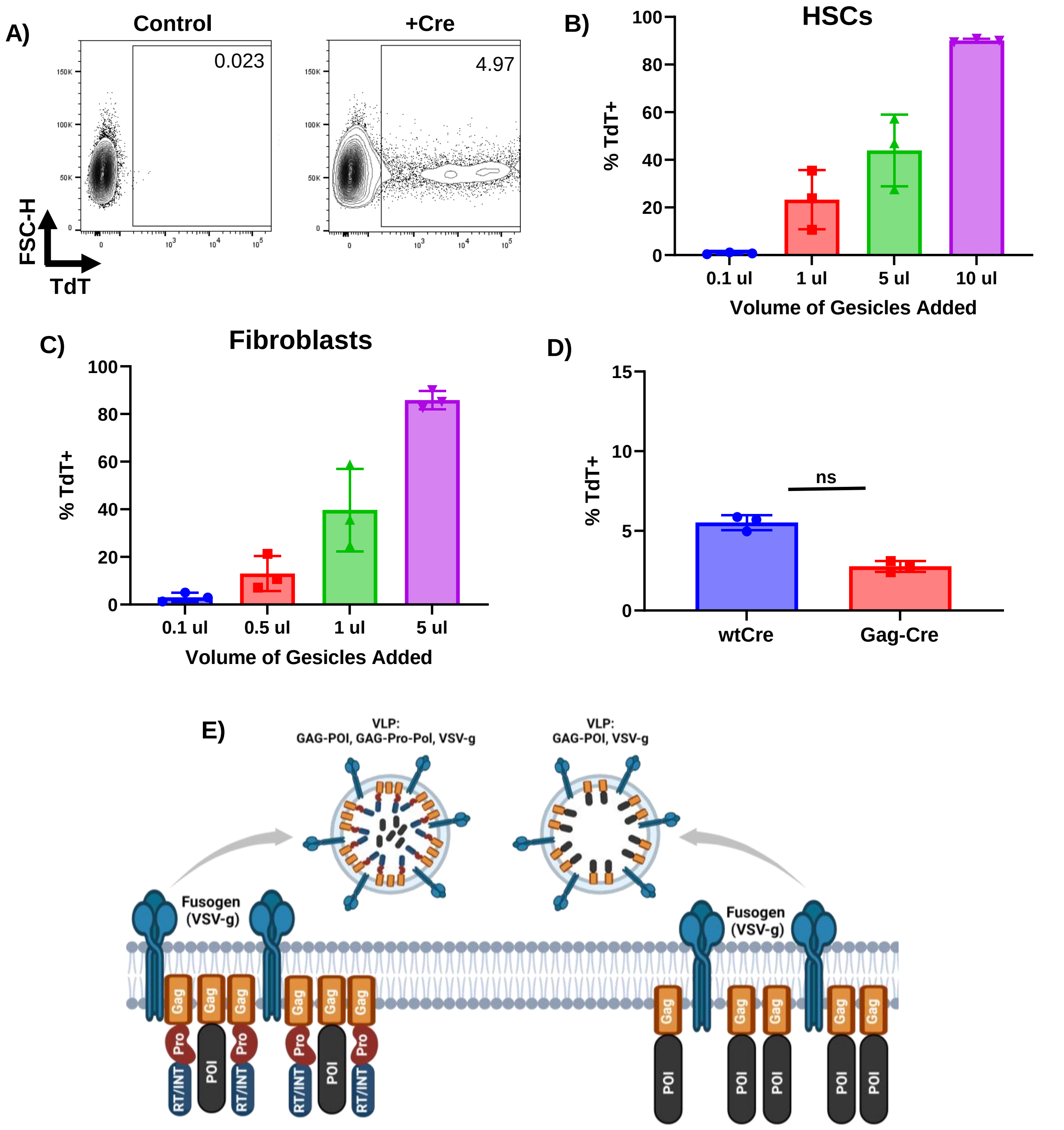


**Supplemental Figure 1: Identification and optimization of a VLP platform to accomplish SPIT. A)** Representative FACS plots detecting Cre recombination in Ai9/Ai14 reporter cells by tdTomato expression using flow cytometry. **B-C)** Frequency of recombination in A14 mouse HSCs and fibroblasts when Gesicles (Takara) are applied to cells across a dose titration (gesicles were pseudotyped with VSV-g) (n=3, mean± s.d). **D)** Bar graph showing the frequency of recombination in Ai9 reporter 293T cells when cells are transfected with plasmids expressing Cre or Gag-Cre (t-test, n=3, mean± s.d). **E)** Schematic of VLP components applied to achieve SPIT and the budding of a VLP from the membrane of a producer cell. The left side shows the production and make up of VLPs produced from expression of GAG-Pro-Pol (Pro, RT/INT), GAG-POI and VSV-g, while the right side shows the make up of VLPs produced from expression of GAG-POI and VSV-g. ns = not significant, POI = Protein of Interest, Pro = MLV Protease, RT/INT = reverse transcriptase/integrase

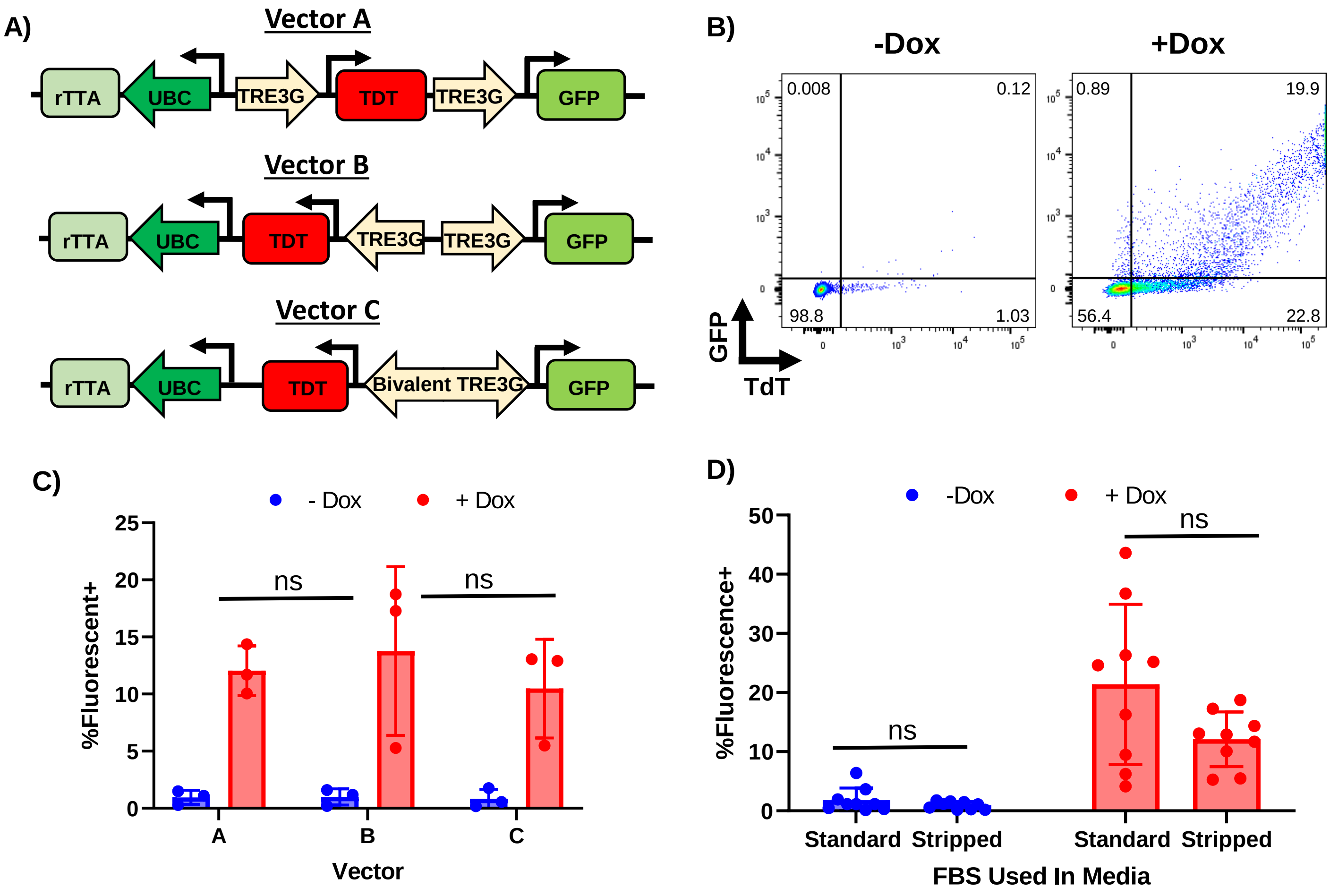


**Supplemental Figure 2: Design & Validation of an All-In-One Doxycycline Inducible Vector. A)** Schematic showing the design and orientation of different all in one doxycycline inducible vectors developed to accomplish doxycycline regulated SPIT. **B)** Representative FACs plots gating for GFP & tdTomato in cells transfected with an all-in-one doxycycline inducible vector when doxycycline was present or absent in the media. **C)** Bar Graph showing the total frequency of cells that were positive for GFP and/or tdTomato expression following transfection with different all in one vectors, when doxycycline was present or absent in the media. No statistically significant differences were found between the expression of transgenes between vectors in the presence or absence of doxycycline (ANOVA, n=3, mean± s.d). Vector A was selected for further use as it had the lowest variance between replicates. **D)** Frequency of fluorescent positive cells following transfection of inducible vectors in the presence or absence of doxycycline, when cells were cultured in media that contained standard FBS or Charcoal Stripped FBS. No statistically significant differences were found between the use of standard FBS or stripped FBS for doxycycline regulation, standard FBS was thus used for inducible SPIT experiments (t-test, n=9, mean± s.d). Dox = Doxycycline.

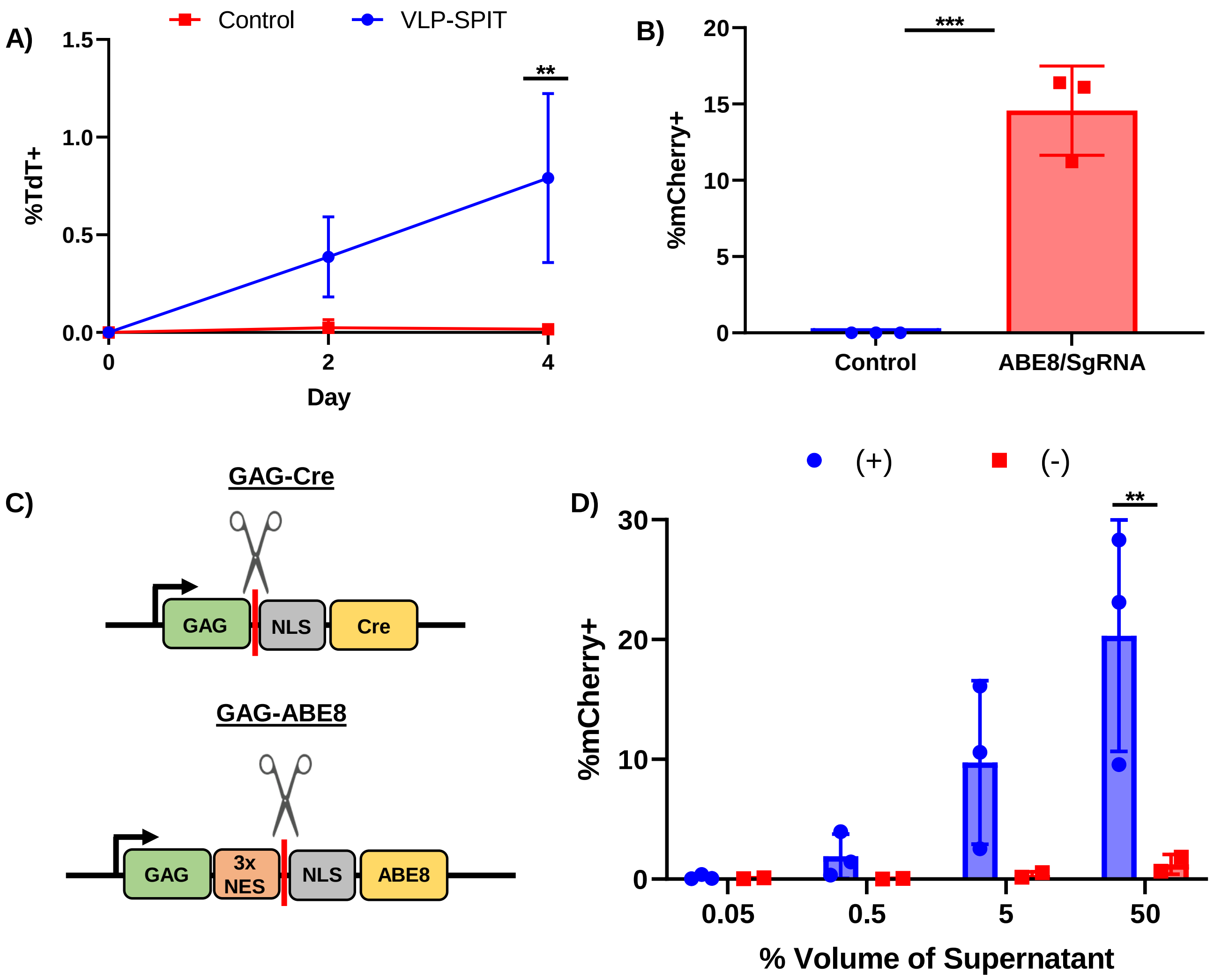


**Supplemental Figure 3: Applying SPIT to Primary Cells and Adapting SPIT for the Delivery of CRISPR-Cas platforms. A)** Line graph showing the frequency of tdTomato+ cells over time when Ai14 mouse fibroblasts are co-cultured with C57BL6/j fibroblasts transfected with VLP-SPIT constructs (GAG-Cre/VSV-g) (ANOVA, n=3, mean± s.d). **B)** Bar Graph showing the frequency of mCherry+ cells when TREE reporter 293T cells are transfected with plasmids encoding an SpCas9 adenine base editor and the TREE sgRNA, compared to untreated TREE cells (t-test, n = 3, mean± s.d). **C)** Schematic showing the difference in the design of Gag-POI vectors that were used to deliver Cre vs ABE8. The scissors and red bar indicate the position at which MLV protease (found within Gag-Pro-Pol) cleaves the POI (Cre or ABE8) from GAG. **D)** Frequency of mCherry+ reporter cells when ABE8 VLPs are produced in the presence (+) or absence (-) of GAG-pro-pol (ANOVA, n=2-3, mean± s.d). Statistical significance determined by ANOVA. ABE8 = SpCas9 Adenine Base Editor version 8. ** = P < 0.005, *** = P < 0.0005.

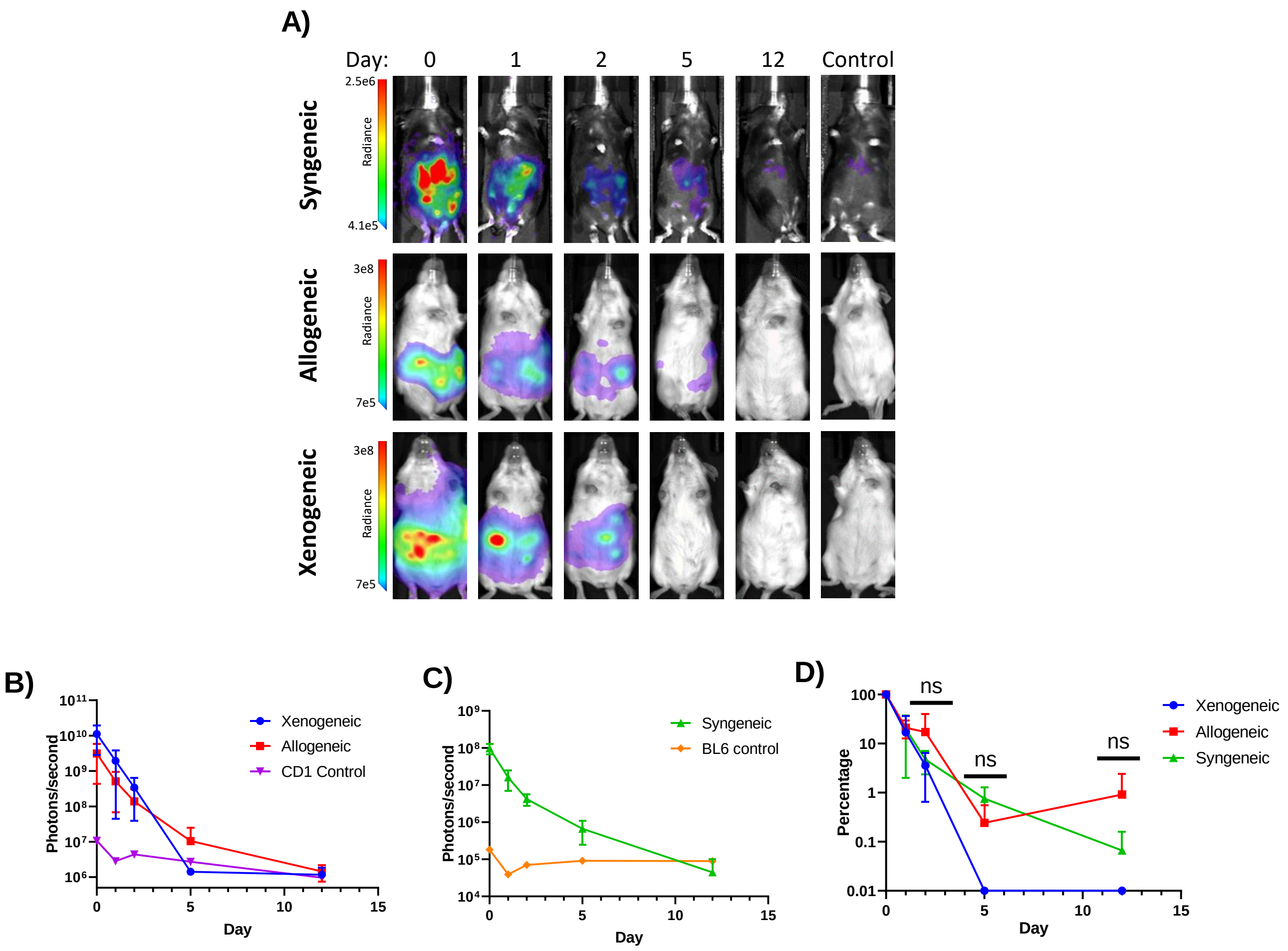


**Supplemental Figure 4: Tracking the persistence of luciferase expressing cells *in vivo*. A)** Representative images from IVIS imaging of mice for luciferase expression after transplanting 2e7 293T cells (xenogeneic) or C57BL6/j mouse embryonic fibroblasts (MEFs) into either CD1 (Allogeneic) or C57BL6/j recipients intraperitoneally. Mice were transfected with luciferase expressing plasmid one day prior to transplantation. **B)** Line graph showing the amount of radiance (photons/s) emitted from CD1 mice transplanted with cells (n = 2-3 experimental, n=1 control, mean± s.d). **C)** Line graph showing the amount of radiance (photons/s) emitted from C57BL6/j (BL6) mice transplanted with cells (n=3 experimental, n=1 control, mean± s.d). **D)** Line graph showing the relative amount of radiance from each experimental condition over time as a percentage of the total radiance detected from mice one hour after transplantation of cells at day 0. No statistically significant differences were found between each group. (ANOVA, n=2-3, mean± s.d)

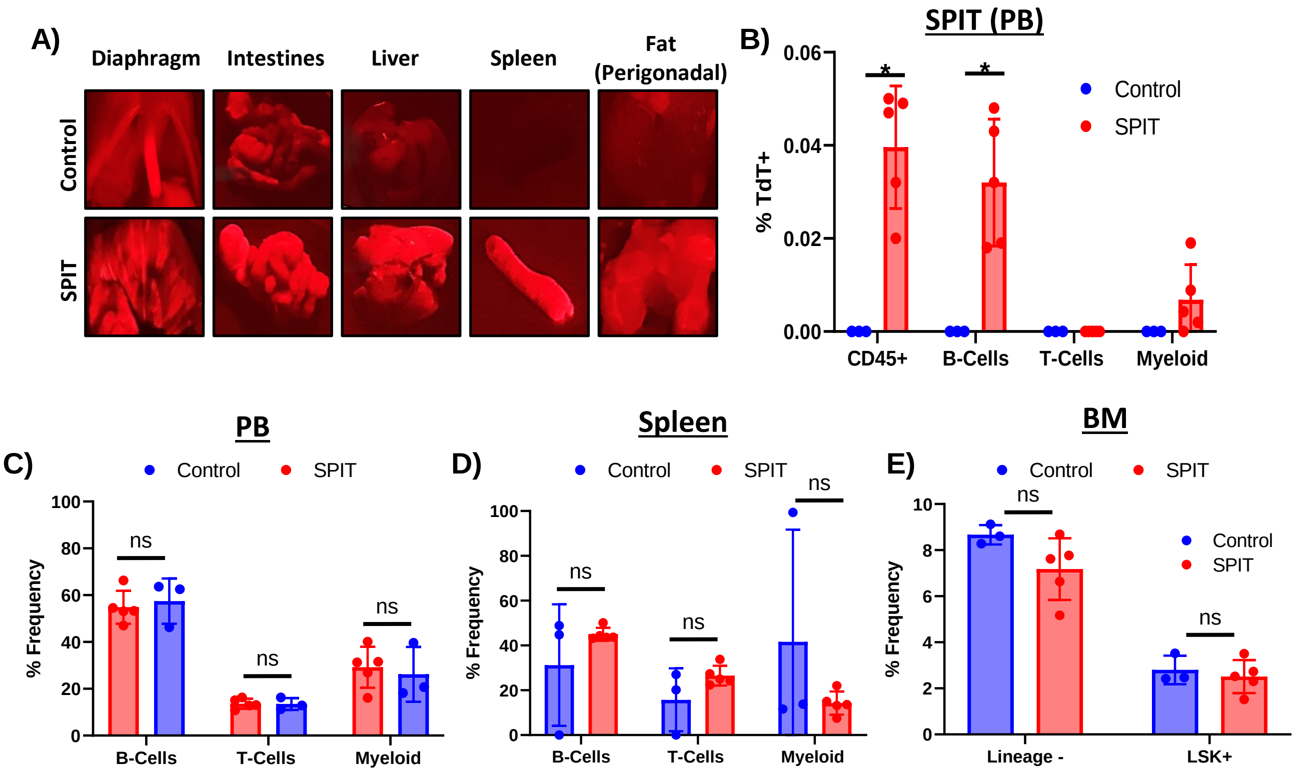


**FIGURE 5: Application of SPIT *in vivo.* A)** Representative images of organs that tdTomato expression could be detected in following the application of SPIT *in vivo* to Ai14 mice compared to organs from untreated control Ai14 mice. A 510-540 nm emission was used to excite tdTomato in organs and a 600 nm filter was used to detect tdTomato signal. **B)** Bar graph showing the frequency of tdTomato expression found in mice that received SPIT compared to control (t-test, n=3-5, mean± s.d). **C-E)** Relative frequency of different cell types found in the peripheral blood **(C)**, spleen **(D)** and bone marrow (BM) **(E)** of mice that received SPIT compared to control mice (t-test, n=3-5, mean± s.d). PB = peripheral blood, BM = Bone marrow, ns = not significant. ** = P < 0.005.

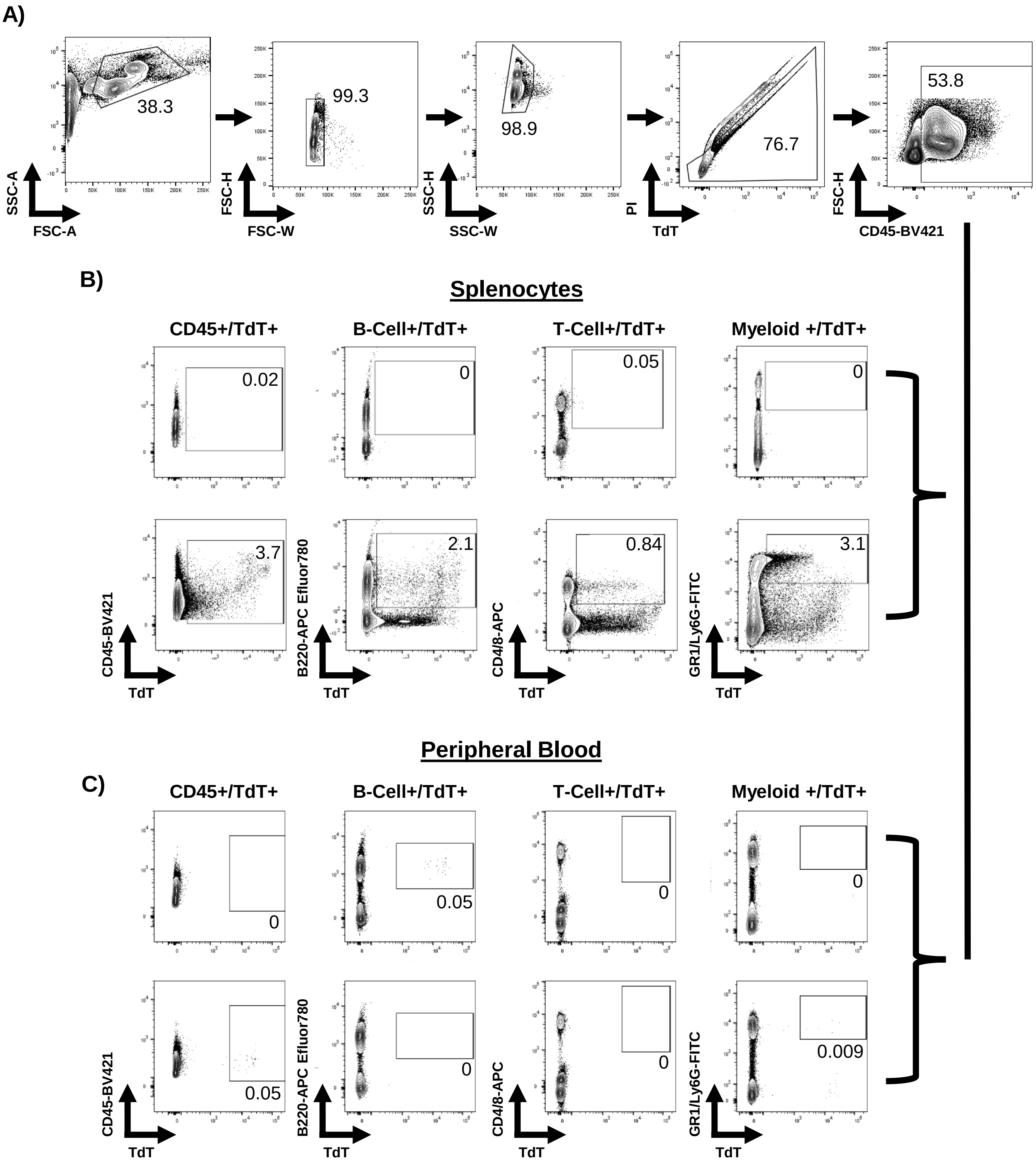


**Supplemental Figure 6: Representative gating from flow cytometry of peripheral blood and splenocytes. A)** Representative images of the gating used on both peripheral blood and spleen for live, CD45+ cells. **B-C)** Representative images of FACs plots from control mice and mice treated with SPIT for detecting tdTomato expression in different cell types of **B)** the spleen and **C)** peripheral blood.

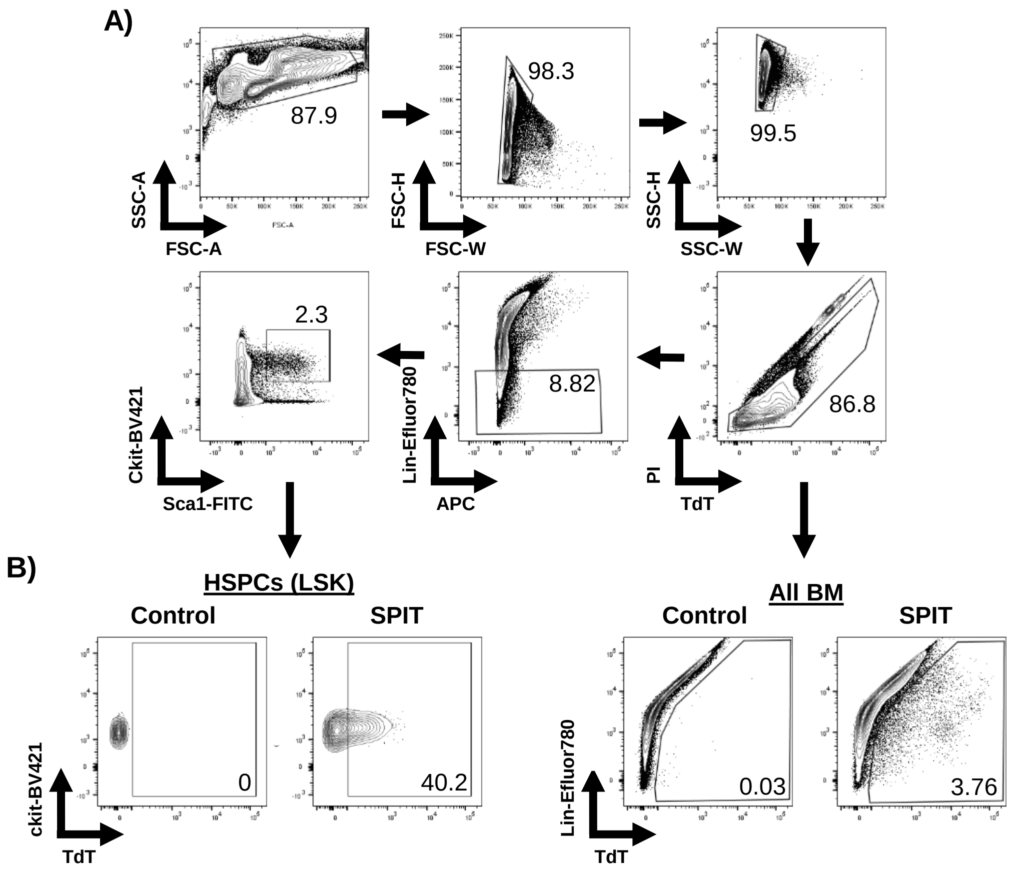


**Supplemental Figure 7: Representative gating from flow cytometry of bone marrow. A)** Representative gating to detect, live, single cells, of different lineages of the bone marrow (by immunophenotyping). **B)** Representative FACS plots looking for tdTomato expression within different subsets of cells within the bone marrow, HSPCs (left) and all cells of the bone marrow (right). Lin = stained with lineage cocktail, refer to methods.
